# Phylodynamics beyond neutrality: The impact of incomplete purifying selection on viral phylogenies and inference

**DOI:** 10.1101/2024.08.28.610037

**Authors:** Katia Koelle, David A. Rasmussen

## Abstract

Viral phylodynamics focuses on using sequence data to make inferences about the population dynamics of viral infectious diseases. These inferences commonly include estimation of the viral growth rate, the reproduction number, and the time of most recent common ancestor. With few exceptions, existing phylodynamic inference approaches assume that all observed and ancestral viral genetic variation is fitness-neutral. This assumption is violated more often than not, with a large body of analyses indicating that fitness varies substantially among genotypes circulating viral populations. Here, we focus specifically on fitness variation arising from deleterious mutations, asking whether incomplete purifying selection of deleterious mutations has the potential to bias phylodynamic inference. We use simulations of an exponentially growing population to explore how incomplete purifying selection distorts tree shape as well as how it shifts the distribution of non-neutral mutations over trees. Consistent with previous results, we find that incomplete purifying selection strongly shapes the distribution of mutations while only weakly impacting tree shape. Despite incomplete purifying selection shifting the distribution of mutations, we find little discernible bias in estimates of the viral growth rate and times of the most recent common ancestor. Our results reassuringly indicate that existing phylodynamic inference approaches may not yield biased epidemiological parameter estimates in the face of incomplete purifying selection, although more work is needed to assess the generalizability of these findings.

## 1 Introduction

This article contributes to a Special Issue commemorating G. Udny Yule’s contribution to the mathematical phylogenetics literature (Yule, 1925). Published 100 years ago in *Philosophical Transactions B*, Yule’s paper ‘A Mathematical Theory of Evolution’ presents a simple generative model for evolutionary diversification. Yule interfaced this model with taxonomic data to estimate model parameters such as speciation rates. Although the details differ considerably, the overall structure of Yule’s approach is reflected in viral phylodynamic analyses today. These analyses commonly involve the specification of a generative model and estimation of model parameters such as growth rates and reproduction numbers (Stadler et al., 2012; Pybus, 2001; Volz, 2012). Model parameters are estimated using reconstructed time-resolved viral phylogenies as the data, with inference approaches nowadays generally integrating over phylogenetic uncertainty (Drummond et al., 2002; Stadler et al., 2013; Volz et al., 2018).

Phylodynamic analyses have also relied on many of the same modeling assumptions as those adopted by Yule in his classic 1925 paper. Most notably, Yule’s model assumes that species-creating mutations are either ‘viable’ or ‘non-viable’ and that all species are equally likely to give rise to the next species. These assumptions lead to the condition of exchangeability being met between species. Similarly, most phylodynamic models ensure exchangeability by assuming all sequence variation is neutral such that all viral lineages share the same fitness regardless of their genotype. While certain models like the multi-type birth-death model allow fitness to vary between lineages based on their genotype, these non-neutral models come at a considerable computational cost even when the number of modeled types is small (Maddison et al., 2007; Stadler and Bonhoeffer, 2013; Barido-Sottani et al., 2020). As a result, the most widely used methods for reconstructing epidemic dynamics from viral sequence data all assume pathogen sequences evolve neutrally.

The assumption that viral mutations are fitness neutral is in direct conflict with a robust set of empirical data. Here, we first briefly review existing analyses that indicate that sublethal deleterious mutations commonly circulate and often contribute substantially to fitness variation between co-circulating viral lineages (section 2). By drawing from the theoretical population genetic literature, we then summarize the impact that incomplete purifying selection is known to have on phylogenetic tree shape as well as on the distribution of mutations on these trees (section 3). We then examine the impact that tree shape distortion alone (section 4) and with non-neutral mutation distributions (section 5) have on viral phylodynamic inference when purifying selection is incomplete. Finally, we discuss our Results in section 6.

## 2 Empirical evidence for incomplete purifying selection in viral populations

Across the diversity of life, most mutations are known to be deleterious. While some of these mutations are lethal deleterious (‘non-viable’), many are sublethal deleterious (Eyre-Walker and Keightley, 2007). In RNA viruses, this pattern also holds true: the overwhelming majority of mutations exact a fitness cost (Sanjuán et al., 2004; Sanjuán, 2010; Visher et al., 2016), with approximately 30% being lethal deleterious and the remaining 70% of mutations being sublethal deleterious. While purifying selection tends to remove deleterious mutations from viral populations over time, selection is generally not strong enough to immediately purge all deleterious mutations. This is especially true if the fitness effects of deleterious mutations and infected population sizes are both small. Furthermore, because viral mutation rates are generally high, the rate at which deleterious mutations enter a viral population can exceed the rate at which purifying selection purges them. Many deleterious mutations can therefore co-circulate in viral populations, and these mutations can cause viral fitness to vary substantially between lineages (Nicolaisen and Desai, 2012; Cvijović et al., 2018).

The imprint of incomplete purifying selection against deleterious mutations can be observed in reconstructed viral phylogenies. Phylogenetic analysis of 143 RNA viruses showed that the ratio of nonsynonymous to synonymous substitution rates on external lineages (i.e., branches leading to sampled tips) was larger than that of internal lineages (Pybus et al., 2007). A simple explanation for this pattern is that purifying selection has had time to filter out lineages carrying deleterious mutations deeper in the tree but not among the more recently sampled lineages.

Incomplete purifying selection may also explain why molecular clock rates estimated from viral sequence data are often found to be time-dependent, with higher rates estimated from samples collected over shorter time scales such as when samples are collected only in the recent past (Ho et al., 2011; Wertheim and Kosakovsky Pond, 2011; Duchêne et al., 2014; Ghafari et al., 2021). Consistent with this explanation, Ghafari et al. (2022) estimated that clock rates are 2 to 4 times higher along external branches relative to internal branches in pandemic H1N1 influenza and SARS-CoV-2 phylogenies. Other studies allowing for branch-specific molecular clocks did not infer such extreme differences in clock rates, with clock rates varying more within internal/external branches than between these groups of branches (Ji et al., 2023). Such extreme differences in clock rates between external and internal branches may therefore only arise when viral sequences are sampled over a relatively short time span, such as early on during an emerging epidemic (Aiewsakun and Katzourakis, 2016; Ghafari et al., 2022).

Several additional studies provide more indirect evidence for incomplete purifying selection in RNA virus populations. One such study focused on predicting clade frequencies of influenza A virus from one season to the next Luksza and Lässig (2014). To predict clade frequency changes, relative fitness values of strains were estimated. For accurate prediction, the authors needed to incorporate fitness costs of amino acid changes that occurred outside of epitope regions (and were thus unlikely to impact antigenicity). Finally, a previous study of ours Koelle and Rasmussen (2015) found that incomplete purifying selection could explain influenza A H3N2’s slender phylogeny and its punctuated antigenic evolution Smith et al. (2004). Among competing antigenic variants, transiently circulating deleterious mutations create additional variation in background fitness in which new antigenic or other beneficial mutations arise, such that beneficial mutations need to occur in higher fitness backgrounds (“jackpot events”) with less deleterious mutation load to displace competing variants. In the absence of circulating deleterious mutations, our simulations instead predict unrealistic levels of viral antigenic diversification in the long-term.

## 3 Brief review of the impact of incomplete purifying selection on phylogenies

Incomplete purifying selection can impact genealogies in two different ways (Williamson and Orive, 2002) (Figure 1). It can distort the shape of a genealogy away from its neutral expectation (Figure 1B), where shape is defined broadly as a tree’s distribution of internal and external branch lengths as well as its balance. It can also impact the distribution of mutations across a genealogy (Figure 1C). Early studies evaluated the impact of purifying selection on tree shape in one-locus, two-allele models (Neuhauser and Krone, 1997; Golding, 1997), finding that purifying selection has only a weak, if at all discernible, impact on tree shape. Later studies transitioned to using simulation-based models that allowed for a larger number of sites and fewer assumptions relating to the fitness costs of deleterious mutations to revisit these conclusions (Williamson and Orive, 2002; Maia et al., 2004). The results of these studies generally corroborated earlier findings with incomplete purifying selection only impacting the shape of genealogies slightly and in certain regions of parameter space. For example, Williamson and Orive (2002) calculated metrics such as the proportion of tree length composed of external branches from simulated genealogies over a range of deleterious mutation fitness costs, finding that this metric was not substantially impacted by incomplete purifying selection. Also using numerical simulations, Maia et al. (2004) found that genealogies derived from contemporaneously sampled individuals were more asymmetrical when mutations exacted a fitness cost, with this effect being most pronounced at large sample size and at intermediate fitness costs of mutations. Consistent with these results, Seger et al. (2010) found that the shapes of genealogies were maximally distorted in terms of branch length proportions when fitness costs of mutations were of intermediate size.

**Figure 1.**
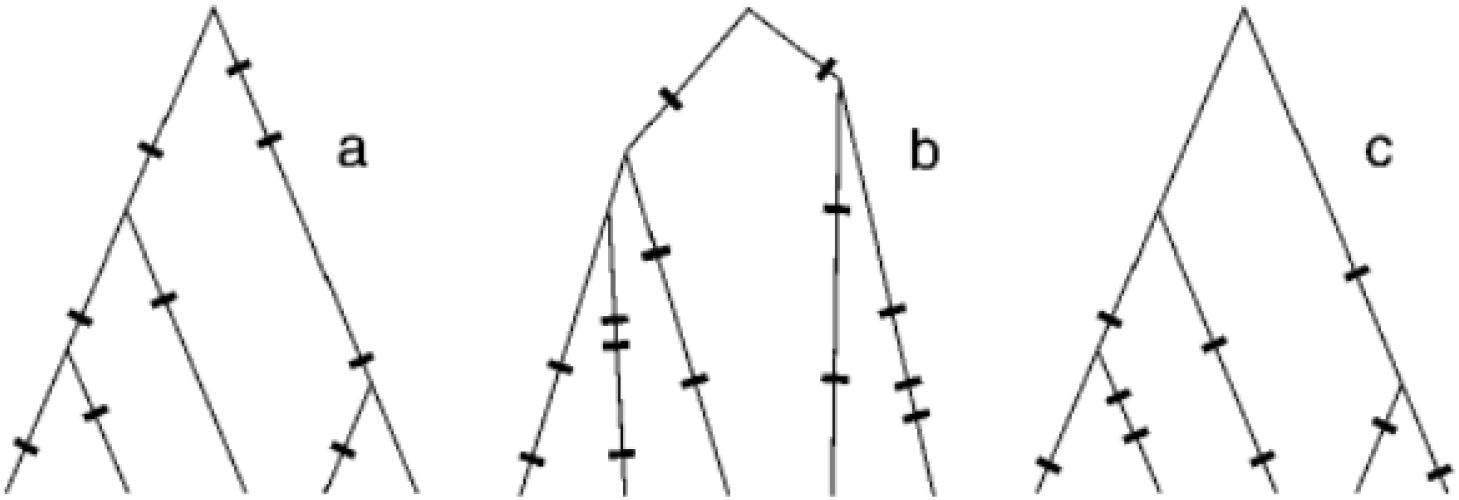
Distinct ways in which purifying selection can impact genealogies, reproduced with permission from (Williamson and Orive, 2002). (A) A neutral genealogy in a constant size population. Both the shape of the tree and the distribution of mutations on the tree is consistent with the neutral expectation. (B) A genealogy that differs only in tree shape from one expected under neutrality. The distribution of mutations is consistent with the neutral expectations. (C) A genealogy that differs only in the distribution of mutations from one expected under neutrality. The tree shape is consistent with the neutral expectation.

Despite not strongly distorting the shapes of genealogies, incomplete purifying selection can nevertheless still strongly shift the temporal distribution of coalescent (branching) events, which could in turn bias demographic inferences under neutral phylodynamic models which typically assume coalescent rates reflect historical populations sizes. Many theoretical models suggest that purifying selection will act to reduce historical effective population sizes, thereby increasing the rate at which lineages coalesce deeper in the past (Seger et al., 2010; O’Fallon et al., 2010; Walczak et al., 2012; Nicolaisen and Desai, 2012). Intuitively, this reduction in effective population size arises because individuals sampled at present have a higher probability of descending from more fit ancestors (Walczak et al., 2012; Neher and Hallatschek, 2013). Thus, the “effective pool” of ancestors that sampled individuals can descend from is reduced relative to neutral expectations where all historical individuals have equal probability of producing sampled descendants. While it has been suggested that the effects of purifying selection on coalescent times can be accounted for by simply rescaling historical effective population sizes (O’Fallon, 2011), others have argued that no such simple rescaling exists (Walczak et al., 2012). Either way, neutral phylodynamic models that ignore how purifying selection can reduce effective population sizes can be prone to biases in demographic inference such as overestimating growth rates or even inferring rapid growth for a constant-sized population (O’Fallon, 2011; Johri et al., 2021).

In contrast to the weak effect that purifying selection seems to have on tree shape, purifying selection can dramatically shift the distribution of mutations on a genealogy.

Specifically, the proportion of mutations that fall on external branches relative to internal branches can be substantially higher in the case of purifying selection than under selective neutrality (Williamson and Orive, 2002), with this proportion increasing at higher strengths of selection *σ*. The clustering of mutations towards the tips of genealogies violates the assumption under neutrality that mutations accumulate at a uniform rate across a tree such that the number of mutations along each branch is Poisson distributed. This can bias branch lengths estimated under the assumption of a constant molecular clock, which in principle could skew coalescent estimates of parameters such as migration rates, effective population sizes *N*_*e*_, and population growth rates (Williamson and Orive, 2002).

## 4 Impact of tree shape distortion on phylodynamic inference

Based on the review above, we first hypothesized that viral phylogenies inferred during an emerging epidemic would not be strongly distorted in shape, and thus, that phylodynamic inference would yield accurate epidemiological estimates when considering only the impact that incomplete purifying selection has on tree shape. To test this hypothesis, we first used SLiM version 4.01 (Haller and Messer, 2019) to forward simulate an exponentially growing viral population across a range of assumed fitness effects for *de novo* mutations. We considered a spillover scenario with a single initially infected individual and used parameters consistent with those for SARS-CoV-2 (Koelle et al., 2022). In the context of an epidemic, births and deaths of individuals represent transmission and recovery events, respectively. We set the birth rate of infected individuals to *λ* = 0.3 per day and the death rate of infected individuals to *δ* = 0.1 per day. These birth and death rates together determine the intrinsic growth rate of the infected population (*r* = *λ − δ* = 0.2 per day), its doubling time (*D* = *ln*(2)*/r* ≈ 3.5 days), as well as the basic reproduction number (*R*_0_ = *λ/δ* = 3).

We let mutations occur at birth, with the number of mutations being Poisson-distributed with mean *m*. We set the per genome, per transmission event mutation rate to *m* = 0.3, again consistent with empirical estimates for SARS-CoV-2 (Park et al., 2023). With a SARS-CoV-2 genome size of 29903 nucleotide sites and a birth rate of *λ* = 0.3 per day, this mutation rate yields a substitution rate of approximately 1.1*×*10^*−*3^ substitutions per site per year under a neutral model of evolution. All mutations within a given simulation had the same fixed fitness cost (*s*_*d*_). We varied fitness effects across simulations and considered *s*_*d*_ values of 0, 0.02, 0.04, 0.08, 0.16, 0.32, and 0.64, with the *s*_*d*_ = 0 simulations serving as the neutral control. We assumed multiplicative fitness effects of mutations, such that a virus with *k* mutations had a fitness of *w* = (1 *−s*_*d*_)^*k*^. We did not consider epistasis of any kind. We forward simulated for *T* = 55 days, and randomly sampled 200 individuals upon recovery from each simulation.

To first assess the extent to which incomplete purifying selection impacts viral tree shape under this epidemic growth scenario, we simulated this forward model under a given *s*_*d*_ parameterization until we obtained 100 simulations with successful viral invasion. Because of demographic stochasticity, the number of infected hosts at *T* = 55 days differed between simulations. From each simulation, we obtained the true viral genealogy of the sampled hosts and used this genealogy to calculate two statistics: the proportion of the total tree length that falls on external branches and the Sackin index of imbalance.

For the neutral forward simulations (*s*_*d*_ = 0), the proportion of the total tree length that falls on external branches was approximately 82.5% (Figure 2A). This proportion is substantially higher than the neutral expectation under a constant-sized population sampled at a single time point, where the expectation is approximately 24% (Fu and Li, 1993). The higher value of this statistic in our simulations is due to exponential growth, which acts to make a phylogeny more ‘star-like’ (Slatkin and Hudson, 1991; Grenfell et al., 2004), with shortened internal branches and lengthened external branches relative to a genealogy from a constant-sized infected host population. Interestingly, the proportion of the total tree length that falls on external branches does not change appreciably even when mutations exact a high fitness cost (Figure 2A). This result is consistent with the findings of Williamson and Orive (2002), which indicated, using this same statistic, that the strength of selection does not appreciably alter branch length distributions in a constant-sized population.

**Figure 2.**
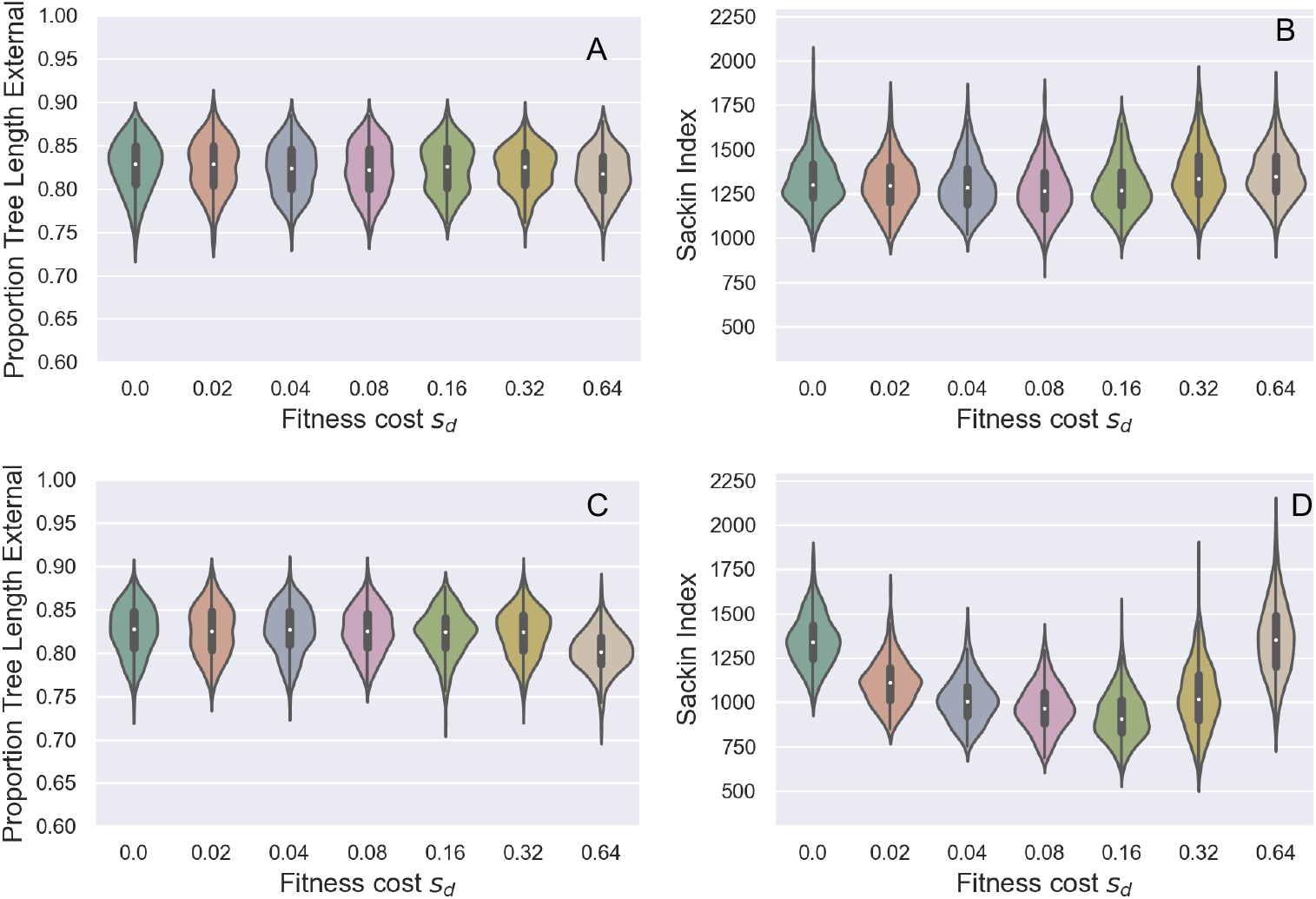
The impact of purifying selection on tree shape across a range of assumed deleterious fitness effects (*s*_*d*_). (A) Proportion of total tree length that falls on external branches. (B) Sackin index of imbalance. In (A) and (B), the mutation rate was set to *m* = 0.30 mutations per genome per transmission. (C) Proportion of tree length that falls on external branches. (D) Sackin index of imbalance. In (C) and (D), the mutation rate was set to *m* = 3.00 mutations per genome per transmission. Violin plots show the distribution of the tree statistics for 100 simulations of the forward model that resulted in viral establishment.

The Sackin index of imbalance is defined as the sum of a tree’s leaves’ depths (Sackin, 1972):

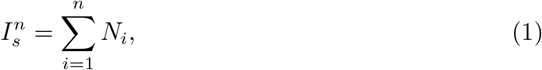

where *N*_*i*_ is the number of internal nodes in the path from leaf *i* to the root and *n* is the total number of leaves in the tree. The value of this index increases with increasing tree imbalance. Here we find that as the fitness cost of deleterious mutations increases, there is no appreciable effect on the Sackin index (Figure 2B). This is perhaps not surprising given previous work that has shown only weak to moderate impact of purifying selection on the average height of a leaf (Maia et al., 2004).

Based on these results, we conclude that tree shape is negligibly impacted by incomplete purifying selection during the early epidemic phase of an acutely-infecting viral pathogens with epidemiological parameters similar to those of SARS-CoV-2. As such, we expect there to be little impact of incomplete purifying selection on phylodynamic inference when only the impact on tree shape is considered. To test this prediction, we randomly selected five genealogies under each fitness cost parameterization. We scattered mutations on these simulated trees, with the number of mutations on a branch being Poisson-distributed with the mean given by the branch length, scaled by the generation interval (1*/λ*) and the mutation rate *m*. We then generated 200 sequences for every genealogy by setting the sequence of the root node to be the SARS-CoV-2 Wuhan-Hu-1 reference isolate (GenBank accession number NC 045512) and assuming an HKY model of sequence evolution with a transition:transversion ratio of 3. We then applied the coalescent exponential model (Griffiths and Tavaré, 1994) implemented in BEAST v1.10.4 (Suchard et al., 2018) to these simulated sequences. From each sequence alignment, we jointly estimated the genealogy, the exponential growth rate *r*, the time of the most recent common ancestor (tMRCA), and the clock rate (Figures 3A-C). Consistent with our expectations, the ‘true’ growth rate of *r* = 0.2 per day was recovered when mutations were neutral (*s*_*d*_ = 0). We were also able to recover the ‘true’ time of the most recent common ancestor (tMRCA) and the ‘true’ clock rate under the neutral model. As expected, when mutations exacted a fitness cost, estimates of the growth rate, tMRCA, and clock rate were still accurate. These results indicate that the slight impact that incomplete purifying selection has on tree shape does not appear to bias phylodynamic inference.

There are several reasons for why tree shape might only be negligibly impacted by purifying selection in our forward simulations. One reason is that deleterious mutations potentially never strongly impact tree shape. This result would be consistent with some of the population genetic work that we reviewed above. However, it may also be that our assumed mutation rate is too low to allow a significant number of deleterious mutations to accumulate over the time span of our simulations. To evaluate this possibility, we resimulated the model with an order-of-magnitude higher mutation rate (*m* = 3.0 mutations per genome per transmission event). At this higher mutation rate, the proportion of total tree length composed of external branches again does not appear to change considerably as the fitness cost of deleterious mutation increases (Figure 2C). However, at this higher mutation rate, the Sackin index of imbalance now appears to depend on the fitness cost of mutations (Figure 2D). Specifically, the Sackin index first decreases with increasing *s*_*d*_ (to approximately 70% of its original value), but then increases again at much higher *s*_*d*_. This result is rather counter-intuitive and inconsistent with (Maia et al., 2004), since we would expect trees to be the most imbalanced at intermediate fitness costs where the most fitness variation would accumulate. Nevertheless, these results indicate that purifying selection has very little overall impact on tree shape regardless of mutation rate.

**Figure 3.**
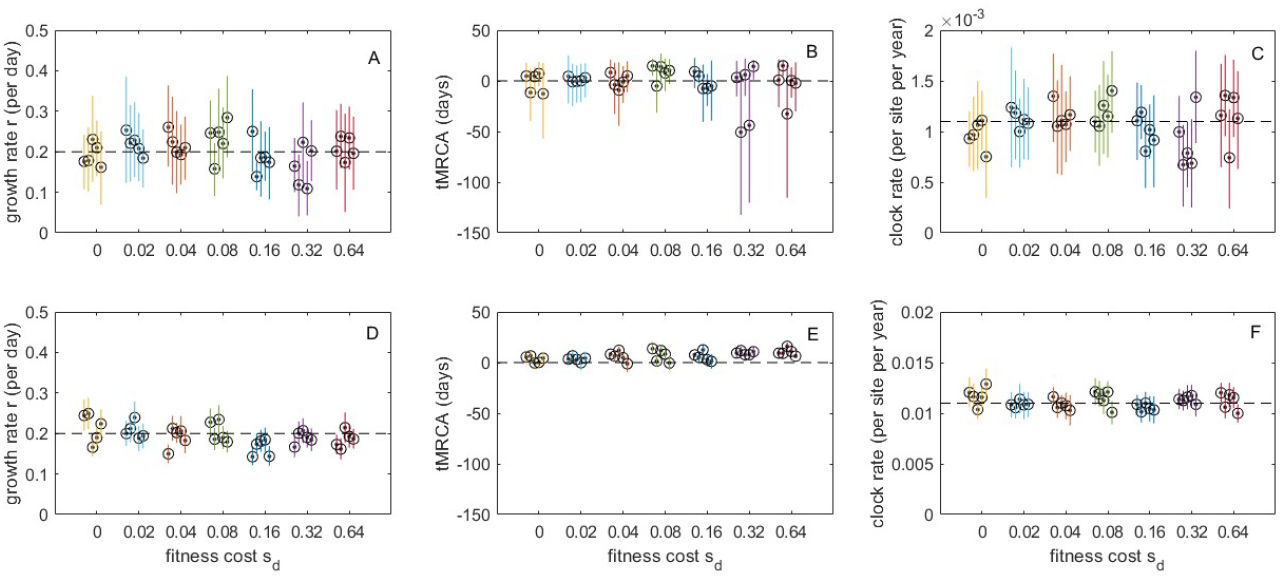
The impact of altered tree shape on phylodynamic inference across a range of mutational fitness costs. (A) Estimated exponential growth rates. (B) Estimated times of the most recent common ancestor. (C) Estimated clock rates. In (A)-(C), mutations were Poisson distributed across branches under the assumption of a mutation rate of *m* = 0.3. (D) Estimated exponential growth rates. (E) Estimated times of the most recent common ancestor. (F) Estimated clock rates. In (D)-(F), mutations were Poisson distributed across branches under the assumption of a mutation rate of *m* = 3.0. At both mutation rates, five replicates are shown for each fitness cost. Circles show mean estimates. Lines show 95% HPD intervals. Dashed black lines show true values. MCMC chain length for all runs was 10^8^, with all effective sample sizes (ESS) exceeding 200 in all runs. Uninformative priors were used for all parameters. For ease of comparison across results, panels (A) and (D) have the same y-axis range and panels (B) and (E) have the same y-axis range.

As before, we scattered mutations on to five randomly selected genealogies for each of the fitness cost simulations according to their branch lengths, scaled by the generation interval and now the higher mutation rate of *m* = 3.0 mutations per genome per transmission. We then again generated 200 sequences for each of these genealogies and applied the coalescent exponential model to these simulated sequences. We were again able to recover the true growth rate of *r* = 0.2 per day under all mutational fitness costs considered (Figure 3D), as well as the tMRCAs (Figure 3E) and the clock rate (Figure 3F). As expected, the credible intervals were tighter under the *m* = 3.0 simulated genealogies than under the *m* = 0.3 simulated genealogies due to less phylogenetic uncertainty under the higher mutation rate. Reassuringly, the impact of incomplete purifying selection on tree balance that is observed at intermediate levels of *s*_*d*_ at the mutation rate of *m* = 3.0 (Figure 2D) does not appear to bias growth rate estimates, tMRCA estimates, or clock rate estimates.

## 5 Impact of mutation distribution on phylodynamic inference

We next consider if the distribution of mutations across viral phylogenies could bias phylodynamic inference in the case of incomplete purifying selection. To do this, we used the same SLiM simulations from section 4, examining this time the distribution of the mutations over the true genealogies. At the mutation rate reflecting empirical values for SARS-CoV-2 (*m* = 0.3 mutations per genome per transmission), incomplete purifying selection had a very strong impact on the distribution of mutations (Figure 4A-C). The proportion of mutations falling on external branches (i.e., singletons, *η*_*e*_*/S* in Williamson and Orive (2002)) increases markedly with the strength of purifying selection (Figure 4A). To see how selection shifts the distribution of mutations independent of distortions in branch lengths, we also compared the proportion of external mutations to the proportion of the tree that consisted of external branches. This ratio grows rapidly with the strength of purifying selection, demonstrating that proportionally more deleterious mutations occur on external branches even after accounting for changing branch lengths (Figure 4B). The shift in the distribution of mutations across the tree also has a major impact on branch-specific clock rates. At the strongest levels of purifying selection (*s*_*d*_ = 0.64) clock rates along external branches are one to two orders of magnitude higher than along internal branches (Figure 4C).

**Figure 4.**
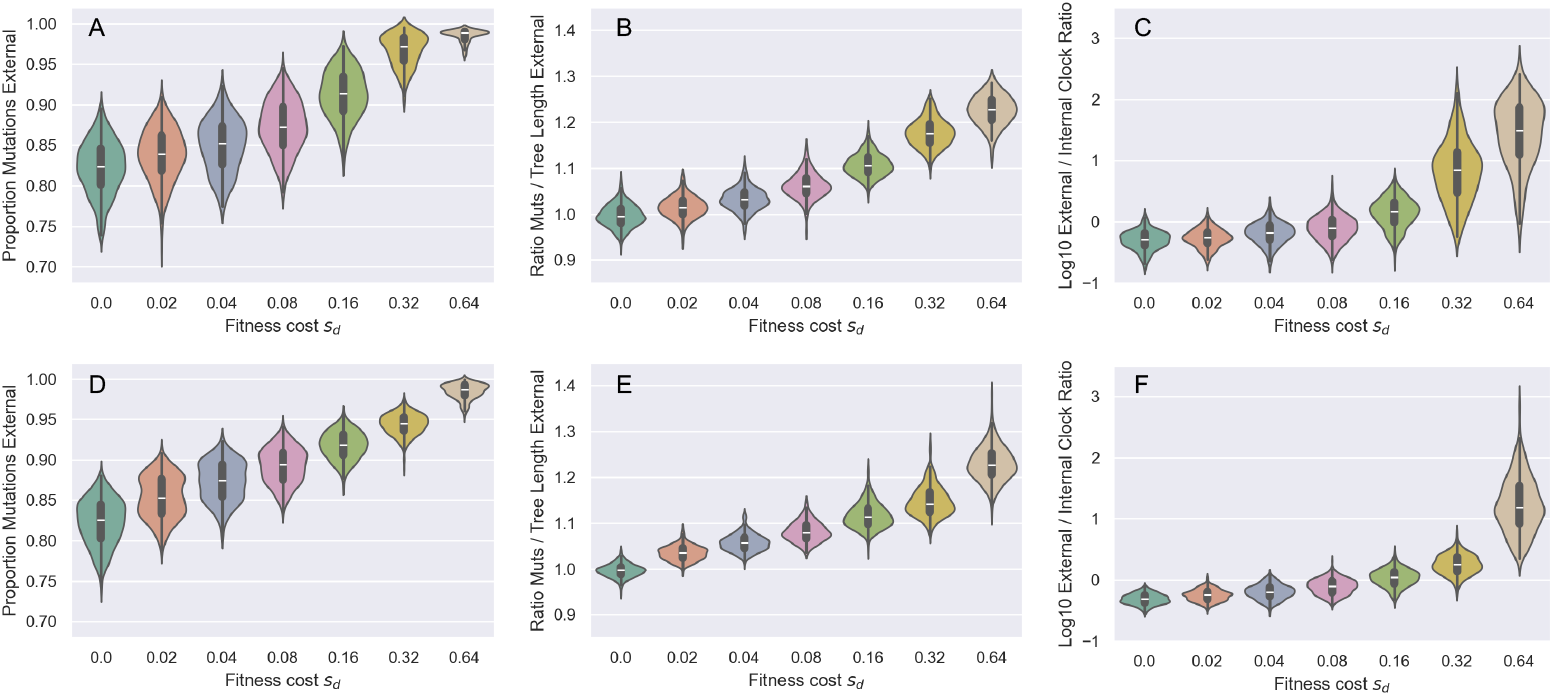
The impact of purifying selection on the distribution of mutations across a range of assumed deleterious fitness effects (*s*_*d*_). (A,D) Proportion of mutations occurring along external branches. (B,E) Ratio of the proportion of external mutations to the proportion of external tree length. (C,F) Ratio of the true external clock rate versus the true internal clock rate. We note that at low values of *s*_*d*_ the (log) ratio is slightly negative. This is because in our simulations mutations only occur at transmission events and we sample individuals upon recovery. The true clock rate is therefore slightly depressed along sampled external lineages. In A-C the mutation rate *m* = 0.30 per genome per transmission event. In D-F, *m* = 3.0 per genome per transmission event. Violin plots show distribution of summary statistics for 100 simulations of the forward model in SLiM.

We repeated these analyses for the simulations parameterized with an order of magnitude higher mutation rate (*m* = 3.0; Figures 4D-F). Again, we found that the proportion of mutations that lie on external branches increases with the fitness cost of deleterious mutations (Figure 4D). The ratio of external mutations to external tree length again increases with increases in the fitness cost of deleterious mutations (Figure 4E), although this ratio appears to increase at a slightly lower rate than it does at the lower mutation rate (Figure 4E vs Figure 4B). Finally, clock rates along external branches are again higher than those on internal branches, although, at a given *s*_*d*_, this ratio is lower at the higher mutation rate than at the lower mutation rate (Figure 4F vs Figure 4C). This is likely because purifying selection can more efficiently remove lineages with an excess of deleterious mutations when mutation rates are high.

Given this observed impact of incomplete purifying selection on the distribution of mutations on our simulated viral phylogenies, we next asked whether phylodynamic inference would still be able to accurately recover epidemiological parameter values. To address this question, we used the same set of true genealogies from section 4, this time simulating sequences for each observed tip based on the true mutations that occurred along each branch. These sequences thus reflect the skewed distribution of mutations on the true genealogies as well as any changes in tree shape that incomplete purifying selection may have left in these genealogies. We then again applied the coalescent exponential model to these simulated sequences to determine whether there would be biases in growth rate or tMRCA estimates. Much to our surprise, we were again able to accurately infer exponential growth rates under both of these mutation rates (Figure 5A, D). At the low mutation rate, we were also able to recover the time of the most recent common ancestor across the range of mutational fitness costs considered (Figure 5B). At the high mutation rate, starting at *s*_*d*_ = 0.08, tMRCA estimates were biased towards more distant past (Figure 5E). At both mutation rates, inferred clock rates were lower the higher the fitness cost (Figures 5C, 5F). These latter findings are unsurprising, given that purifying selection decreases the number of observed substitutions along the trees, particularly on internal branches. The biases in tMRCA estimates (Figure 5B) may be a result of these lower clock rates: the tMRCA is most likely estimated to occur in the more distant past because we underestimate the clock rate, requiring more time to have elapsed between the tMRCA and the tips to explain the observed genetic distances between sequences.

**Figure 5.**
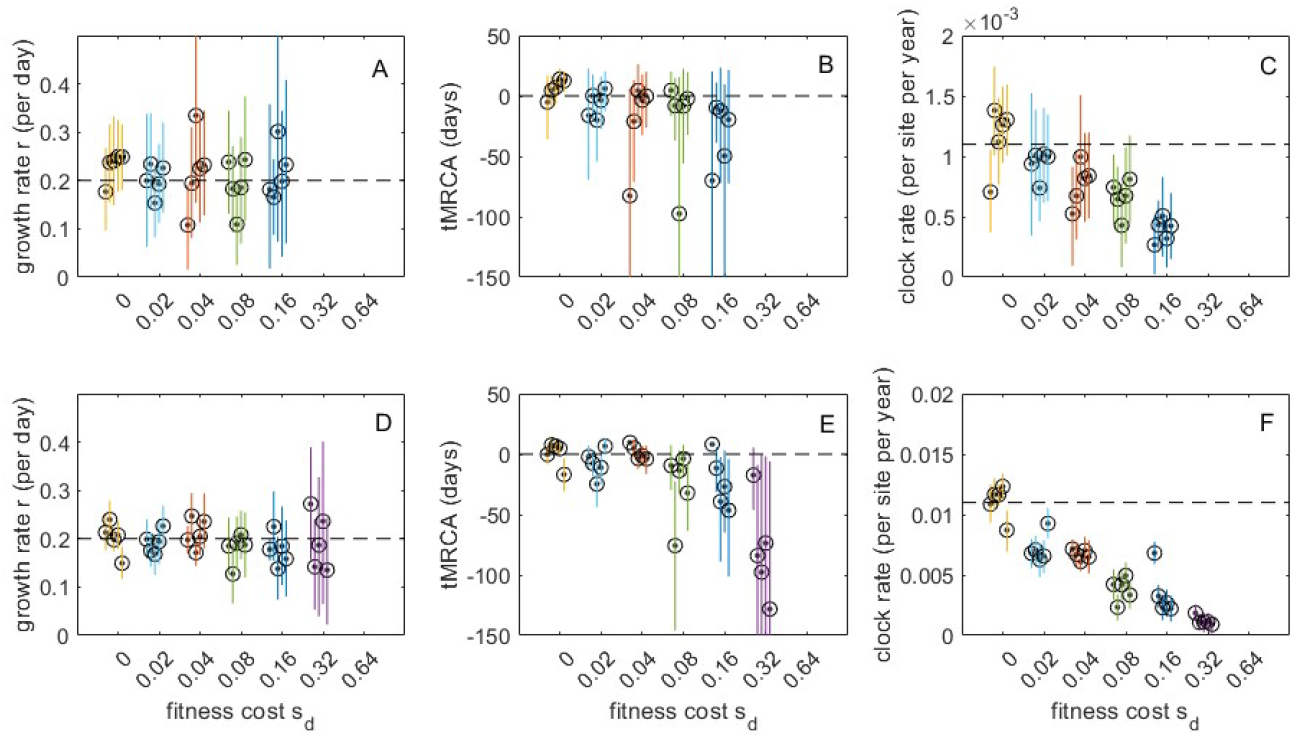
The impact of altered mutation distribution (jointly with tree shape) on phylodynamic inference across a range of assumed deleterious fitness costs. (A,D) Estimated exponential growth rates. (B,E) Estimated times of the most recent common ancestor. (C,F) Estimated clock rates. In (A-C), the mutation rate *m* = 0.3. In (D-F), the mutation rate *m* = 3.0. At both mutation rates, five replicates are shown for each fitness cost. As in Figure 3, circles show mean estimates and lines show 95% HPD intervals. Dashed black lines show true values. With the exception of *m* = 3.0 and *s*_*d*_ = 0.32, MCMC chain length for all runs was 10^9^, with all effective sample sizes (ESS) exceeding 200 in all runs. Inference on sequences from *m* = 3.0 and *s*_*d*_ = 0.32 simulations combined 2-3 log files, each with an MCMC chain length of 10^9^ such that effective sample sizes (ESS) exceeded 200 in all 5 replicates. Inference was not possible under a parameterization of *m* = 0.3 and *s*_*d*_ = 0.32, under a parameterization of *m* = 0.3 and *s*_*d*_ = 0.64, or under a parameterization of *m* = 3.0 and *s*_*d*_ = 0.64 due to insufficient amounts of genetic variation between sampled individuals. Uninformative priors were used for all parameters. For ease of comparison across results, panels (A) and (D) have the same y-axis range and panels (B) and (E) have the same y-axis range.

## 6 Sensitivity to sampling times

The impact of incomplete purifying selection on viral phylogenies and phylodynamic inference may be dependent on when sampling occurs. For example, we may have only been able to accurately estimate epidemic growth rates in the face of strong purifying selection because we sampled viral lineages continuously through time, including early on when few deleterious mutations are segregating. We therefore may have been able to accurately infer growth rates from the distribution of coalescent times among early sampled lineages before purifying selection begins to distort the coalescent process. Likewise, purifying selection may have the greatest impact on estimated clock rates when samples are only collected over a short period of time in the recent past (Ho et al., 2011; Duchêne et al., 2014; Aiewsakun and Katzourakis, 2016; Ghafari et al., 2022). We therefore explored how sensitive our results are to different sampling schemes in our *m* = 0.3 mutation rate simulations. We conducted this sensitivity analysis by varying the time period over which samples were collected. As a point of comparison to our previous simulations where we sampled continuously over the full *T* = 55 days of exponential growth, we considered cases where the sampling period was constrained to the final 20 days and then the final 5 days of the simulation.

As before, incomplete purifying selection has little impact on tree shape in terms of the proportion of external tree length or the Sackin index, regardless of the sampling period (Figure 6A-B). External lineages become longer when sampling is restricted to the more recent past, but this is expected for any exponentially growing population where the population size is larger in the more recent past resulting in more recently sampled lineages coalescing at a slower rate than lineages sampled at earlier time points. In contrast to tree shape, sampling in the more recent past tends to exacerbate the impact of purifying selection on the distribution of mutations. The proportion of singleton mutations falling on external branches increases the later sampling occurs regardless of *s*_*d*_ value (Figure 6C), although at very high *s*_*d*_ values nearly all mutations occur along external lineages regardless of when sampling occurs. As a result of this, sampling in the more recent past has a larger impact on the proportion of external tree length than on the proportion of external mutations when the fitness costs of mutations are very large (Figure 6A versus 6C). Thus the ratio of the proportion of external mutations to the proportion of external tree length can actually decrease as sampling occurs later at large *s*_*d*_ values (Figure 6D). Nevertheless, clock rates remain higher along external lineages relative to internal lineages when sampling occurs in the more recent past (Figure 6E), consistent with earlier empirical observations (Ghafari et al., 2022).

**Figure 6.**
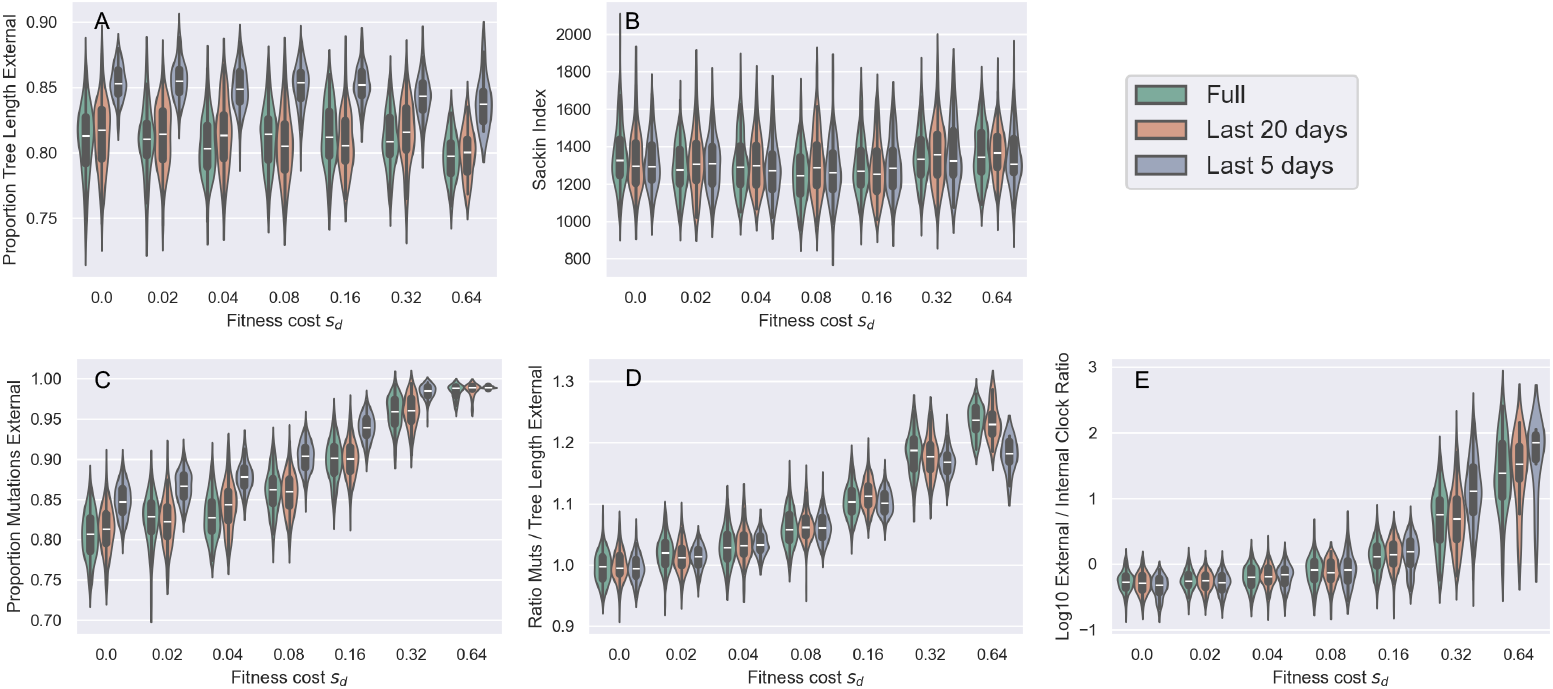
The impact of purifying selection on tree shape and the distribution of mutations when sampling occurs over different time periods. Violin plots are colored according whether samples were collected over the full epidemic (green), the final 20 days (orange) or the final 5 days (blue). (A) Proportion of tree length composed of external branches. (B) Sackin index of imbalance. (C) Proportion of mutations occurring along external branches. (D) Ratio of the proportion of external mutations to the proportion of external tree length. (E) Ratio of the true external clock rate versus the true internal clock rate. The mutation rate was *m* = 0.30 per genome per transmission in all simulations. In all simulations, we sampled 200 individuals at random from the set of individuals that recovered over the sampling period. Because of this, and the exponential growth of the infected population, there are always a larger number of individuals sampled toward the end of the time period considered.

To determine whether phylodynamic inference could still recover the true epidemiological parameter values when sampling of individuals was constrained to a short period of time, we again generated sets of sequences from the simulations with 200 individuals sampled over the final 5 days of the epidemic between *T* = 50 days and *T* = 55 days. As in Figure 5, the sequences we generated reflected the mutation distributions of the true genealogies. We again attempted to estimate the growth rate, tMRCA, and clock rate under the coalescent exponential model. Again, to our surprise, we were able to accurately recover the growth rate *r*, the tMRCA, and the true clock rate (Figure 7). Although our 95% HPD intervals generally contained the true values of these parameters, these credible intervals were (as expected) much wider when individuals were sampled over a five day period compared to when individuals were sampled over the full time period of the epidemic) (Figure 7 versus Figure 5A-C). Due to these wide credible intervals, it is also difficult to determine whether sampling over a highly constrained period of time indeed biases phylodynamic inference.

**Figure 7.**
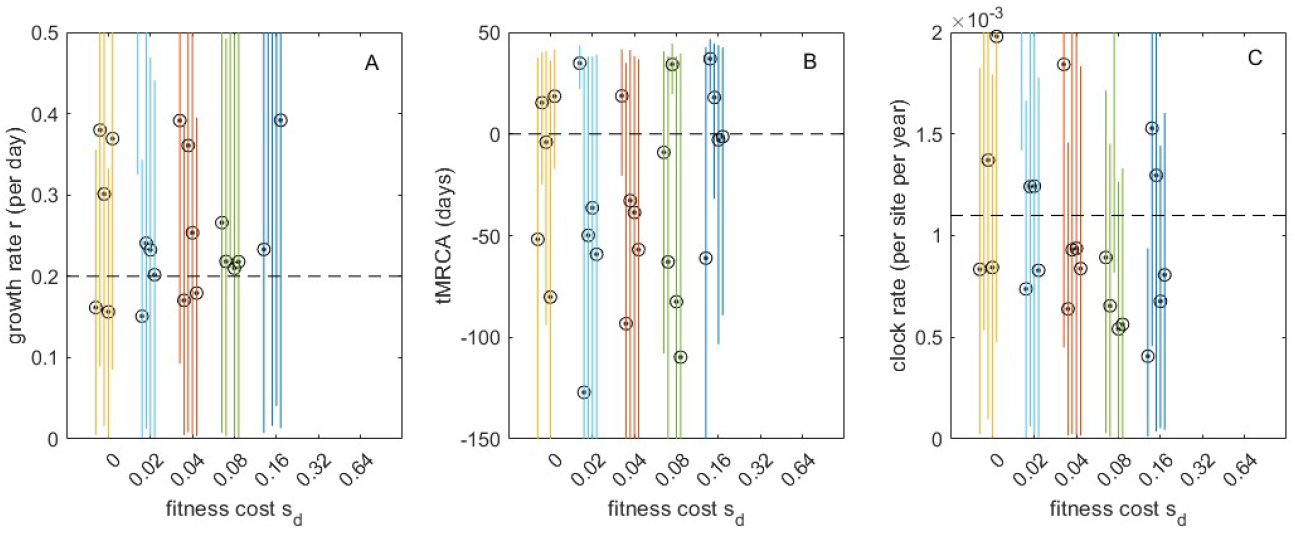
The impact of incomplete purifying selection on phylodynamic inference across a range of deleterious fitness costs, for simulations with a mutation rate of *m* = 0.3 and sampling only from *T* = 50 through *T* = 55 days. (A) Estimated growth rate. (B) Estimated time of the most recent common ancestor. (C) Estimated clock rate. All MCMC chain lengths were 10^9^ or combined across multiple log files to obtain ESS values exceeding 200. Uninformative priors were used for all estimated parameters.

## 7 Discussion

Phylodynamic inference methods, including both coalescent-based and birth-death approaches, generally assume that all observed and ancestral viral genetic variation is fitness-neutral. Here, we first briefly reviewed studies showing that this assumption is violated for the majority of studied viral pathogens, including those circulating in humans. These studies have generally found that purifying selection occurs in viral populations, although it is incomplete, resulting in transient circulation of sublethal deleterious mutations. We then reviewed existing findings from the population genetic literature that have assessed the impact of incomplete purifying selection on tree shape and mutation distribution. Based on these results, we anticipated that tree shape would only be slightly distorted from its neutral expectation in viral populations undergoing epidemic expansion. In contrast, we further anticipated that the distribution of mutations across these genealogies would be starkly impacted by incomplete purifying selection, with a shift in mutations towards the external branches of trees and away from internal branches. As a result of these expectations, we hypothesized that phylodynamic inference could be biased as a result of changes in the mutation distribution away from that expected under a neutrally-evolving, exponentially growing population.

Consistent with the first set of our predictions, we found that, under a parameterization that reflected values estimated for SARS-CoV-2 during its early 2020 viral expansion, tree shape was not substantially impacted by incomplete purifying selection across a broad range of considered mutational fitness costs (Figure 2A-B). Phylodynamic inference applied to simulated sequence data generated by exclusively considering impacts on tree shape was able to successfully recover the true epidemiological parameters, as expected (Figures 3A-C). Again consistent with our predictions, we then found that the distribution of mutations across trees indeed shifted substantially as mutations exacted an increasing fitness cost (Figure 4A-C). Surprisingly, however, phylodynamic inference was able to nevertheless recover the true growth rate and the true tMRCA. Clock rates, as expected, were lower at higher mutational fitness costs, due to purifying selection. At an order-of-magnitude higher mutation rate as well as at a reduced sampling period of 5 days, phylodynamic inference was still able to recover the true epidemiological parameters of interest, again, much to our surprise (Figures 5D, 5E; Figures 7A, 7B).

Our analyses involved a limited number of scenarios and many assumptions. First, we largely considered a single parameterization with a growth rate of *r* = 0.2 per day and a mutation rate of *m* = 0.3 mutations per genome per transmission event, although we did explore a considerably higher mutation rate scenario (*m* = 3.0). We also considered only an epidemic invasion scenario, where a total of 200 individuals were sampled within 55 days of the start of the epidemic, although we also explored sensitivity to sampling over a more constrained period of time. As such, we are not certain of the generalizability of our results. It could be that under a different parameterization or under a different sampling scheme, that incomplete purifying selection could bias parameter estimates from phylodynamic inference. One possible suggestion is therefore for inference to first be done under an assumption of neutrality, followed by an analysis similar to those presented here at the estimated parameter values to determine whether one might expect a bias in parameter estimates. Finally, we restricted ourselves to inference using the coalescent exponential model to examine the impact of incomplete purifying selection on phylodynamic inference. We anticipate that inference based on the birth-death model would yield similar results, but we have not performed these analyses. In sum, our (albeit limited) analyses indicate that phylodynamic inference is not biased by the circulation of deleterious mutations that fall between ‘viable’ and ‘non-viable’. That parameter estimates from phylodynamic inference align as closely with the true epidemiological parameter values of our simulations as they do, despite pervasive incomplete purifying selection, is, as Yule would put it ‘in fact better than one has any right to expect’.

